# One step forward, two steps back: transcriptional advancements and fermentation phenomena in *Actinobacillus succinogenes* 130Z

**DOI:** 10.1101/2020.12.31.424933

**Authors:** Dianna S. Long, Cheryl M. Immethun, Lisbeth Vallecilla-Yepez, Mark R. Wilkins, Rajib Saha

## Abstract

Within the field of bioproduction, non-model organisms offer promise as bio-platform candidates. Non-model organisms can possess natural abilities to consume complex feedstocks, produce industrially useful chemicals, and withstand extreme environments that can be ideal for product extraction. However, non-model organisms also come with unique challenges due to lack of characterization. As a consequence, developing synthetic biology tools, predicting growth behavior, and building computational models can be difficult. There have been many advancements that have improved work with non-model organisms to address broad limitations, however each organism can come with unique surprises. Here we share our work in the non-model bacterium *Actinobacillus succinognes* 130Z, which includes both advancements in synthetic biology toolkit development and pitfalls in unpredictable fermentation behaviors. To develop a synthetic biology “tool kit” for *A. succinogenes*, information gleaned from a growth study and antibiotic screening was used to characterize 22 promoters which demonstrated a 260-fold range of fluorescence protein expression. The strongest of the promoters was incorporated into an inducible system for tunable gene control in *A. succinogenes* using the promoter for the *lac* operon as a template. This system flaunted a 481-fold range of expression and no significant basal expression. These findings were accompanied by unexpected changes in fermentation products characterized by a loss of succinic acid and increase in lactic acid after approximately 10 months in the lab. Contamination and mutation were ruled out as causes and further testing is needed to elucidate the driving factors. The significance of this work is to share tools developed in *A. succinogenes* while simultaneously serving as a cautionary tale. In sharing our findings, we seek to provide necessary information for further development of *A. succinogenes* as a platform for bioproduction of succinic acid. Additionally, we hope to illustrate the importance of diligent and long-term observation when working with non-model bacteria.

## 1. Introduction

Recent research endeavors have turned to generating useful chemicals from biological platforms as an environmentally responsible alternative to non-sustainable sources (1, 2). Bioproduction of industrially important chemicals can utilize organic and renewable feedstocks as nutrient-sources for microbial fermentation using metabolically engineered strains for optimized production. Examples of bioproduction success stories include the production of artemisinin (an anti-malaria drug) from engineered yeast (3) and hydrogen from engineered *E. coli* (4). Non-model organisms are becoming increasingly interesting bioproduction platforms as they would expand the range of metabolic capabilities potentially harnessed for bioproduction purposes. Specific characteristics that would make an organism a good biological platform include native abilities to degrade sugar polymers, utilize renewable feedstocks to produce biochemicals of interest, and grow in challenging environments (5). These unique characteristics that can be found in non-model microbes go hand-in-hand with unique challenges. The limited characterization of non-model organisms can raise issues when utilizing synthetic biology tools in predictable ways, elucidating effective metabolic engineering strategies, and understanding complex regulatory behaviors. Advances in computational tools to harness omics data and synthetic biology have made it possible to begin development of non-model organisms as bioproduction platforms. In fact, a recent review highlights many success stories of how challenges of working with non-model organisms have been overcome to unlock their unique potential (6). Examples include the identification and incorporation in centromeric regions to solve the problem of low plasmid maintenance in the lipogenic and unconventional yeast *Yarrowia lipolytica* (7) and the modification of CRISPR/Cas9 plasmid system to reduce problematic recombination events in non-model actinobacteria producers of diverse natural products (8) among many others. These innovative solutions can serve as inspiration for work in other non-model organisms, such as *Actinobacillus succinogenes* 130Z.

*A. succinogenes* 130Z is a Gram-negative, biofilm-forming, capnophilic, anaerobic and non-model bacterium identified as a potential bioproduction platform for succinic acid (hereafter SA) (9) and could also be developed for other products such as itaconic acid or fumarate. Here we target production of SA; an organic acid that can serve as a precursor for many chemicals used in the production of various commodities, including biodegradable plastics, active pharmaceutical agents, and textiles (10-12). It has been predicted that bioproduction of SA from complex sugar sources could become the primary mode of production, eventually replacing current unsustainable methods that rely on declining petroleum sources (13-15). SA bioproduction is supported by *A. succinogenes’* ability to utilize both C5 and C6 sugars derived from cellulosic biomass (9, 16) and its unique metabolic pathway which includes a truncated TCA cycle resulting in naturally high production of SA (17) without demonstrating product inhibition (16). This bacterium is a biosafety level 1 organism meaning it could be readily incorporated into any industrial facility. To this date, *A. succinogenes*-driven SA production has reached a yield of 94% (w/w) from glucose (16, 18) yet has a theoretical yield of 121% (w/w) from glucose (19). It has also been demonstrated to grow robustly on corn stover hydrolysate which contains chemicals that can inhibit microbial growth (20). This indicates that this bacterium could be an efficient SA producer through its ability to utilize the carbon in hydrolysate without requiring extensive preprocessing. It has been noted that growth condition optimization is not sufficient to obtain maximum SA levels (17), therefore, increasing SA production further will require other strategies such as metabolic engineering using synthetic biology tools, few of which exist for this non-model organism. Previous studies have shown strategies employing endogenous promoters (21) and gene-knock out methods (21-23), but as of yet, exogenous promoters have not been tested or characterized in *A. succinogenes* and no specific inducible promoter has been designed for this bacterium. It is well-known that development of promoters, specifically inducible promoters that can be turned on and off, is one of the easiest and most effective ways to control gene expression (24). Hence, a wider range of available tools would allow for further fine-tuning of *A. succinogenes*’ metabolism for maximizing SA production.

To this end, here we share a case study of both advancements and challenges of working with *A. succinogenes* for SA production. Several steps were taken prior to engineering *A. succinogenes* to increase SA production, including performing small-scale growth studies, identifying effective selection antibiotics, and characterizing and developing synthetic biology tools. We show characterization of 22 constitutive promoters using green fluorescent protein and a flavin-binding fluorescent protein demonstrating a 260-fold range of expression from the weakest to strongest promoter. Additionally, we present characterization of the commonly used inducible *lac* system from *E. coli* and our development of a novel inducible system demonstrating a 481-fold dynamic range. While the progress toward a synthetic biology toolkit for *A. succinogenes* is an important development, we also find SA production being lost over time in the working stock of *A. succinogenes*. This unexpected fermentation shift is yet to <u>be</u>overcome, however, we believe that both the progress and the challenges shared here will aid in future development of *A. succinogenes* as a more stable and efficient producer of SA. This would benefit not only the stakeholders (i.e., bioprocessing industry and related agricultural markets) but also open new doors for bioproduction efforts using other non-model microbes.

## 2. Results and Discussion

### 2.1 Growth curve

Identifying *A. succinogenes’* growth phases (lag, exponential, and stationary) is important as some molecular biology methods (e.g. electroporation (25)) require their application within certain growth phases. In literature, growth conditions have been described in 500 mL flasks (26), 500 mL Duran bottles (27), bioreactors (18, 21, 26, 28), and test tubes (29); however, there has yet to be a reported growth curve showing the growth phases that are important for small scale engineering studies. A growth curve for *A. succinogenes* was generated by fitting a logistic model to OD_600_ measurements taken at 30-minute intervals over the course of a 10-hour growth period. Results demonstrated a 2-hour lag phase, followed by a 6-hour exponential growth phase after which the cells entered stationary phase (Supplementary Figure S1). Early exponential phase, a key point for transformation of the bacterium via electroporation (25), was determined to be between hour 2 and 4 and at 0.4-0.6 OD_600_.

### 2.2 Antibiotic Screening

Another crucial aspect for developing synthetic biology tools is effective selection antibiotics. Screening of standard antibiotics is needed to provide options that enforce plasmid maintenance for tool testing. Cultures of *A. succinogenes* were grown in the presence of kanamycin (50 µg/mL), tetracycline (10 µg/mL), ampicillin (100 µg/mL), gentamicin (15 µg/mL), spectinomycin (50 µg/mL), or chloramphenicol (34 µg/mL). Antibiotic concentrations were selected within the ATCC recommended concentration range for plasmid maintenance in bacteria containing mid-range plasmid copy number. Previous work with *A. succinogenes* shuttle vectors demonstrated low to medium copy number (30), therefore concentrations within the guidelines for mid-range copy numbers should be sufficient. OD_600_ was measured at 2, 4, 6, and 24 hours. Figure 1 shows the efficacy of each antibiotic displayed as percent growth repression (Materials and Methods). Although spectinomycin and ampicillin both eventually inhibit growth of this fast-growing bacterium equivalently to the other tested antibiotics, neither took effect until after two hours. The delayed response to spectinomycin may be due to slow uptake, which is possible when using aminoglycoside class antibiotics in anaerobic conditions (31). Similarly, the slow response to ampicillin may be due to difficulty passing through the cell wall, which can be seen in Gram-negative bacteria (32). Although wild-type *A. succinogenes* cells were eventually killed, using ampicillin for selection posed a problem when selecting successfully transformed colonies of this biofilm-forming bacterium. Resistance to ampicillin is achieved by excretion of β-lactamase which breaks open the antibiotic’s β-lactam ring (33). It is possible that cells containing the plasmid could create an environment for the non-transformed cells to keep growing by excreting the enzyme into the growth medium, therefore making selection difficult. Based on these findings, kanamycin, tetracycline, chloramphenicol, and gentamicin are recommended as selection markers in *A. succinogenes*.

**Figure 1:**
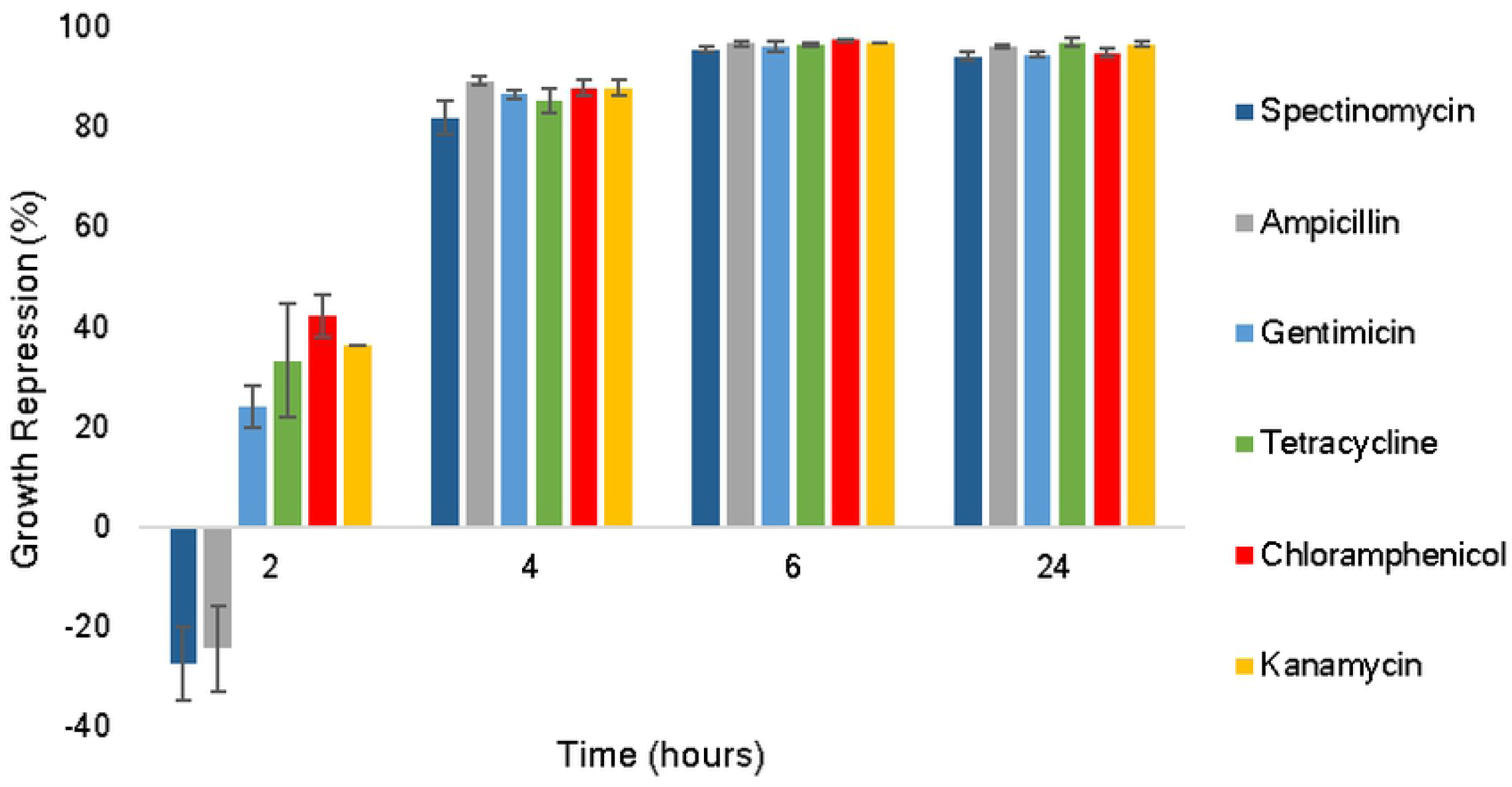
Antibiotic screening in liquid media with wild type *A. succinogenes* shown as percent repression of normal growth. Error bars represent one standard deviation.

### 2.3 Constitutive promoter library

To enable control of transcription in *A. succinogenes*, 22 constitutive (always expressing) promoters were characterized. The set of promoters included P_*lac*_, a promoter native to *E. coli* (34); P_*pcka*_, a promoter native to *A. succinogenes* (21); and the family of Anderson promoters, synthetic promoters that were developed in *E. coli* (35). The Anderson promoter library and the Lac operon’s promoter were chosen since they have been shown to function well in multiple bacteria (36-40). A major goal of this project was to investigate the use of synthetic biology tools that would be less likely to demonstrate cross talk with native genetic components in *A. succinogenes* (38). Characterization of these orthogonal tools within *A. succinogenes* was necessary as promoters often do not behave the same across different species, which can be seen in the cyanobacterium *Synechocystis* sp. 6803 (38). For example, in *Synechocystis*, promoter BBa_J23112 was stronger than BBa_J23100 whereas in *E. coli*, BBa_J23100 produced the strongest expression. Furthermore, P_*lac*_ was not inducible with IPTG in the cyanobacterium. As shown in Figure 2A, each promoter was inserted into the plasmid SSBIO-AS001 upstream of the modified jellyfish *A. victoria* green fluorescence protein gene (*gFPuv*) (41). Transformation of the plasmids into *A. succinogenes* created strains sAS100 – sAS122 (Supplementary Table S5) and normalized expression (Materials and Methods) is shown in Figure 2B. To compare the relative expression of the Anderson promoter library in *A. succinogenes* with activity in *E. coli*, average expression values in *A. succinogenes* were divided by the highest expressing promoter (BBa-J23100) thus setting the maximum expression to 1. These values were compared to reported relative expression values in *E. coli* (35), which were calculated in the same way, as can be seen in Figure 2C. Findings demonstrated that maximal and minimal expressing promoters were the same in both bacteria however, relative expression was not equivalent across the entire promoter set. For instance, BBa_J23119 and BBa_J23100 were expressed similarly in *E. coli* while there was a drastic difference in expression of the two promoters in *A. succinogenes*, revealing unique sensitivities between the two bacteria to sequence variations at different locations within the promoters. A look at how the expression varies within the promoter set reveals a pattern that is evident in both *E. coli* and *A. succinogenes*. A guanine instead of a thymine at the position −12 appeared to hinder expression. This location falls within the −10 hexamer region and the decreased expression may be due to a lower affinity of polymerase binding. Aside from that one consistency, there is not a clear pattern of how the promoter sequence is tied to expression changes, thus reiterating the importance host-specific characterization. A second reporter gene, flavin-binding fluorescent protein (42), hereafter *fbfp*, was inserted in place of *gFPuv* and was tested under the control of a mid-range Anderson promoter (BBa_J23111) and the native promoter (P_*pcka*_). FbFP was used because the protein can fold in anaerobic conditions (42) unlike GFPuv which requires oxygen (43). This allowed for fluorescent measurements to be taken immediately after culturing and removed any variation that may have been introduced by the aeration process used when measuring GFPuv expression. Results showed that BBa_J23111 expressed both GFPuv and FbFP approximately 4 times stronger than the native promoter P_*pcka*_ (4.49 and 4.29 respectively), demonstrating consistently stronger expression with the synthetic promoter. Finally, to determine the range of expression across all tested promoters in *A. succinogenes*, promoters showing no expression were discarded and the remaining 15 were compared to the lowest expressing promoter (BBa_J23105) showing a relative range of 260-fold. The Anderson promoters demonstrate a significantly greater range of expression (up to 2,547-fold) in *E. coli* (35) (verification shown in Supplementary Figure S2). A possible explanation is that the Anderson promoters were designed from *E. coli*’s consensus sequence (labeled as BBa_J23119 in the Anderson promoter set). This means that minor changes were made to the −10 and −35 hexamer regions of the optimum promoter sequence for transcription within *E. coli*. In contrast, *A. succinogenes*’ consensus sequence has not been determined. While *E. coli* and *A. succinogenes* share many characteristics, there may be variation within the replication machinery, such as differences in sigma factors (44), which could account for the lower range of expression observed in *A. succinogenes*.

**Figure 2.**
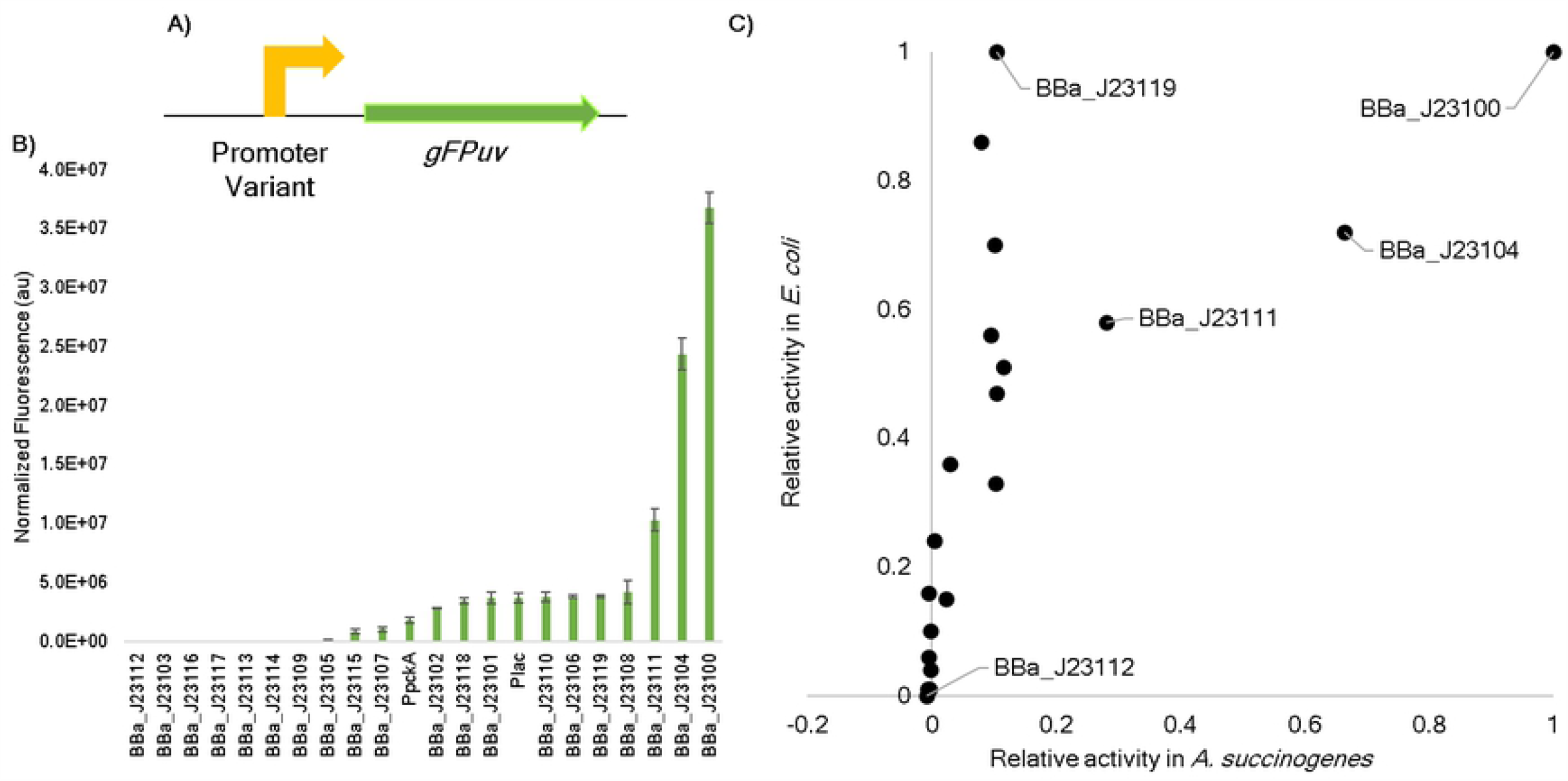
Characterization of 22 constitutive promoters in *Actinobacillus succinogenes* 130Z. (A) Schematic of SSBIO-SA001 expression system. (B) GFPuv expression normalized by absorbance and background control (Materials and Methods). Error bars represent one standard deviation. (C) A comparison of relative expression of the Anderson promoters in *A. succinogenes* and *E. coli*. For both bacteria, relative expression was calculated by dividing all values by normalized expression from the strongest promoter (BBa_J23100).

### 2.4 Characterization of the inducible P_*lac*_ promoter

While constitutive promoters are useful tools for setting constant gene expression rates, metabolic engineering often requires promoters that can respond to external signals (45). The *lac* operon from *E. coli* (34) is an inducible system that includes the promoter P_*lac*_ flanked by operator regions that bind to the repressor protein, LacI. When LacI is bound, transcription is turned off via steric hinderance; however, an inducer molecule, isopropyl β-D-1-thiogalactopyranoside (hereafter IPTG), can bind to LacI and prevent repression of transcription. Therefore, varying levels of IPTG can tune expression of genes under control of P_*lac*._ To characterize this inducible system within *A. succinogenes*, the repressor protein’s gene, *lacI*, and its native promoter were included with all the same components as SSBIO-AS001, creating SSBIO-AS003 (Figure 3A). Transforming *A. succinogenes* with the plasmid created strain sAS124. Performance of sAS124 induced at 0 and 5 mM IPTG was compared to both wild type *A. succinogenes* and strain sAS120 containing the constitutive system with P_lac_. Growth was compared across IPTG concentrations and was shown to be consistent (Supplementary Figure S3). As can be seen in figure 3B, expression in sSA124 at 0 mM IPTG was not statistically different than wild type background fluorescence and expression in sSA124 at 5 mM IPTG was not statistically different than sAS120. Cultures were grown at various concentrations of IPTG within the range of 0 to 5 mM and expression was measured. Visualization on a logarithmic scale showed a graded response (Figure 3C). While the change in expression from the deactivated to activated state of the inducible system was only 90-fold, the on-state matched the expression level of constitutive P_lac_, demonstrating complete induction. Additionally, the system did not show leakiness, as no GFPuv expression was observed in the absence of IPTG. While the range of control is limited by the relatively low maximum level of expression of P_*lac*_ within *A. succinogenes* when compared to other constitutive promoters (see Figure 2B), the binary on and off states make this system promising for development within *A. succinogenes*.

**Figure 3.**
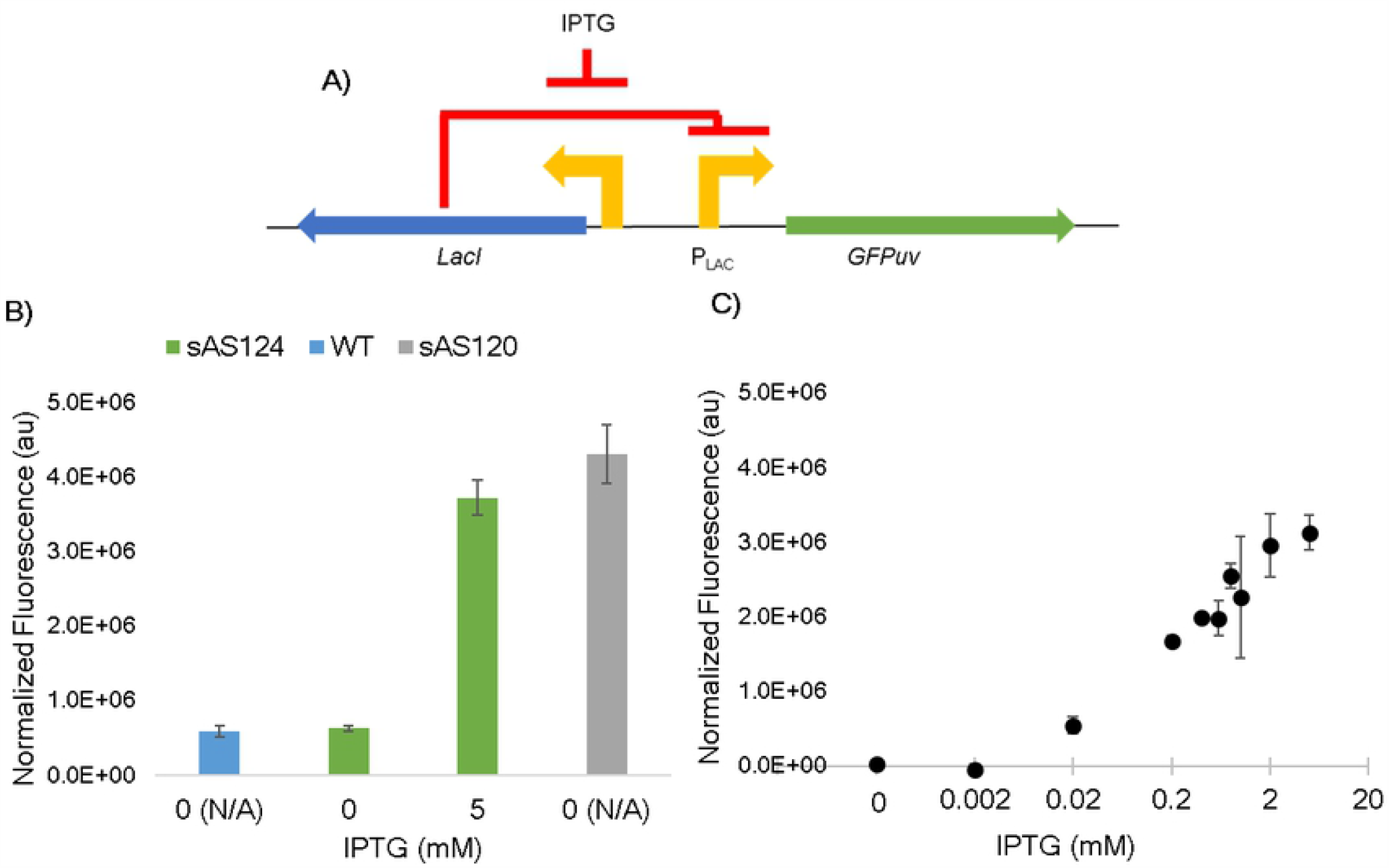
Characterization of strain sAS124 containing SSBIO-AS003 using GFPuv as the reporter protein. A) Schematic of the expression system used in SSBIO-AS003. B) Comparison of on and off states of sAS124 at 0 and 5 mM IPTG with wild type *A. succinogenes* and strain sAS120. Values are normalized by absorbance (Materials and Methods). C) Induction of sAS124 at IPTG levels ranging from 0 to 5 mM. Values are normalized by absorbance and wild-type (Materials and Methods). All error bars represent one standard deviation.

### 2.5 Development of a stronger inducible promoter: p100i

To create an inducible system for *A. succinogenes* with a larger range of expression, the strongest constitutive promoter (BBa_J23100) was added to SSBIO-AS003 in place of the core P_lac,_ sequence creating SSBIO-AS004. In the constitutive system, BBa_J23100 expressed GFPuv ∼10 times stronger than P_lac_ and was predicted to set a higher maximum level of expression for the inducible system. The design strategy followed work in cyanobacterium (45, 46) and can be seen in Figure 4A. Due to the differences in length of the core promoter between P_lac_ (36 bp) and BBa_J23100 (35 bp), each possible nucleotide was inserted on the 5’ end of the −35 region of BBa_J23100 (hereafter called position −36). Position −36 was chosen to keep the spacing between the operator sites O1 and O3 as well as between the −10 region and the transcription start site equivalent for the new promoter and P_lac_. As can be seen in Figure 4B, there was significant variation among the four versions (strains sAS125-sAS128). A cytosine allowed for the greatest expression level whereas guanine and adenine showed decreased expression and thymine showed the least expression. Expression tests in *E. coli* revealed the same pattern, suggesting that the single nucleotide was crucial for some aspect of transcription. Sequences upstream of the −35 hexamer can have various regulatory effects due to interactions with transcription factors (47). For example, the Cyclical-AMP receptor protein (CRP), which enhances polymerase binding, has a binding site in the *lac* system upstream of the −35 hexamer and improves transcription efficiency. It is possible that variations at the −36 site may be changing the binding affinity for CRP. Further exploration of *A. succinogenes* replication machinery will need to be conducted to elucidate the variation caused by the −36 site residues.

**Figure 4.**
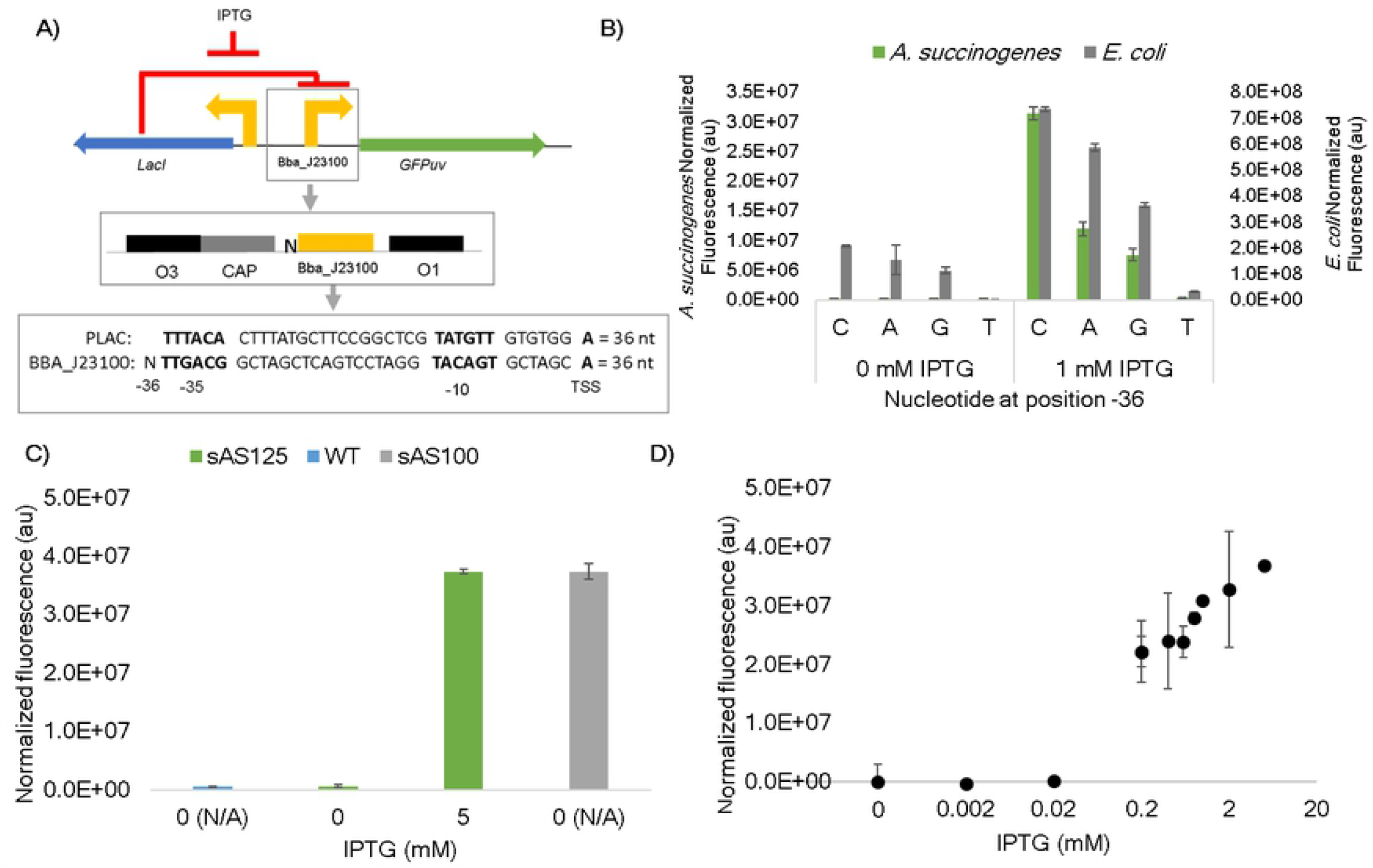
Characterization of strain sAS125 containing SSBIO-AS004 using GFPuv as the reporter protein. A) Schematic of the expression system used in SSBIO-AS004 showing strategy of where to include the extra nucleotide to make size and spacing equivalent. B) Comparison of expression variation due to the −36 nucleotide in both *A. succinogenes* and *E. coli* normalized by absorbance and respective wild type (Materials and Methods) at 0 and 1 mM IPTG. C) Comparison of on and off states of sAS125 at 0 and 5 mM IPTG with wild type *A. succinogenes* and strain sAS100. Fluorescence values are normalized by absorbance (Materials and Methods). D) Induction of sAS125 at IPTG levels ranging from 0.002 to 5 mM. Fluorescence values are normalized by absorbance and wild-type. All error bars represent one standard deviation.

Performance of sAS125, containing the cytosine −36 residue, and which is hereafter labeled p100i, induced at 0 and 5 mM IPTG was compared to both wild type *A. succinogenes* and strain sAS100 containing the constitutive system with BBa_J23100. As can be seen in figure 4C, expression in sSA125 at 0 mM IPTG was not different than wild type and expression in sSA125 at 5 mM IPTG was not different than sAS100. These findings confirmed that p100i was demonstrating a non-leaky off-state and complete induction. Cultures were grown at various concentrations of IPTG within the range of 0 to 5 mM. Visualization on a logarithmic scale showed a graded response (Figure 4D) and 481-fold dynamic range. The non-leaky nature of this inducible system is unique as even *E. coli* shows leakiness of basal expression (48), making this a very exciting development towards *A. succinogenes* metabolic engineering.

### 2.6 Loss of Succinic Acid Production

At this stage, we sought to test the effect of the presence of the plasmids on the production of SA and discovered a decrease in production not only present in the transformed strains, but also in the wild type used for comparison. This loss was in relation to SA measurements that had been taken in the wild type *A. succinogenes* at the very beginning of the project, approximately 10 months before the synthetic biology tools had been constructed and characterized and also in relation to a brand new strain purchased after the loss was realized (Figure 5a). In addition to the loss in SA production, we also observed an increase in lactic acid. This unsettling finding prompted further investigation. The initial hypotheses were either contamination or mutation causing the decrease in SA production, both of which were tested by sequence comparisons between producing (hereafter SA(+)) and non-producing (SA(-)) strains. Using primers for 16S rRNA, sequencing of the SA(-) strain revealed a 99% match with *A. succinogenes*’ reference genome (GCA_000017245.1), making contamination an unlikely cause. Whole genome comparisons between producing and non-producing strains revealed no major mutations, but one small, 5 nucleotide deletion in SA(-) at position 731146 in an intergenic region and a single nucleotide polymorphism (A in SA(+) and reference, G in SA(-)) at position 1004969 at the 5’ end of ASUC_RS04870 (Figure 5b). Investigation into these mutations reveal the SNP has been previously shown to have no effect on SA production (49). While the deletion would have to be investigated to completely rule out mutation as the cause for the loss of SA production, such a minor difference between the two genomes in an intergenic region suggests there may be a better explanation.

**Figure 5.**
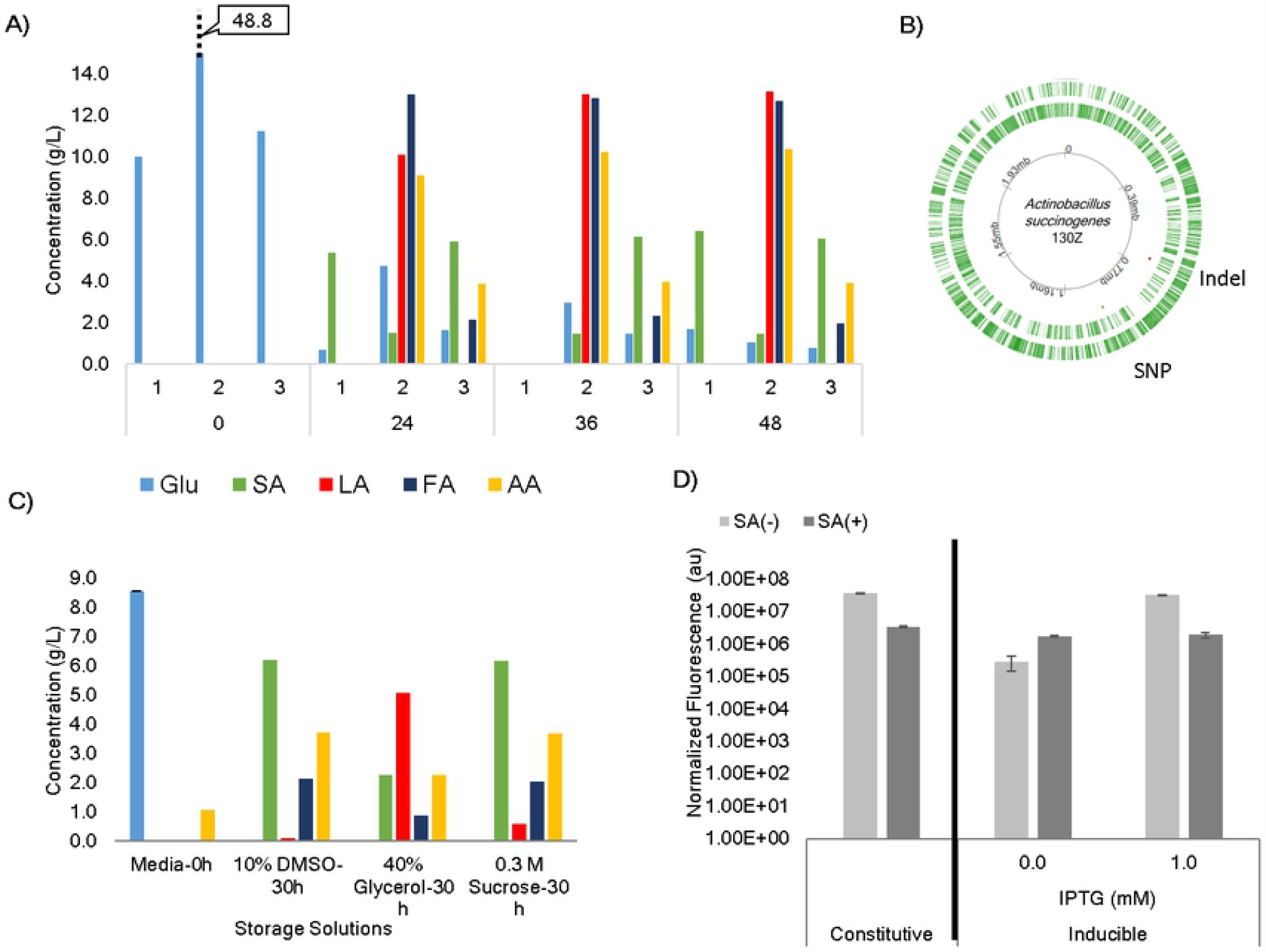
Findings related to succinic acid production loss in *A. succinogenes*. A) Comparison of organic acid fermentation profiles between *A. succiongenes* working stock at the beginning of use in the lab (1), ten months later (2), and a newly purchased strain (3). B) *A. succinogenes* genome map indicating positions of the indel mutation (loss of 5 nts) and SNP mutation (A->G) shown to be different in the SA(-) strain. C) Fermentation profiles of *A. succinogenes* after storage in DMSO, sucrose, and glycerol after 10 freeze/thaw cycles. D) GFP expression data in SA(+) and SA(-) strains under the control of BBa_J23100 and p100i. Glucose (Glu), succinic acid (SA), lactic acid (LA), formic acid (FA), acetic acid (AA).

Since the loss of SA production occurred after several months, one factor may be storage stress. Cells had been stored in a 50/50 mixture with 40% glycerol at −80°C. To see if a different cryo-protectant would prevent fermentation shifts, aliquots of a newly purchased SA(+) strain was stored in various cryo-protectants including the intracellular protectants glycerol (40%) and DMSO (10%), and the extracellular protectant sucrose (0.3M). Aliquots were then subjected to 10 freeze-thaw cycles to mimic use in the lab and SA fermentation at 30 hours was measured. The findings are shown in Figure 5c and indicate there is some difference in optimality of cryo-protectant used, with both DMSO and sucrose showing better protection than glycerol. Cells stored in glycerol show the pattern of decreased SA and increased lactic acid that we saw in the old strains used in the lab. However, there was not a complete loss of SA production and so this was not completely sufficient to explain the shift in fermentation profile seen after 10 months.

For further comparisons between SA(+) and SA(-) strains, the constitutive and inducible promoters, BBa_J23100 and p100i, were transformed into a new SA(+) strain and expression tests were run. This revealed that the differences between SA(+) and SA(-) strains extended to how the synthetic biology tools worked. As seen in Figure 5d, the tools were much less effective in the SA(+) strain with the inducible promoter p100i losing inducibility. This finding adds to the picture of what is happening. Not only are there changes in overflow metabolism, but also changes in the expression of genes on a plasmid. This suggests that gene regulators were playing a role in the phenotypes of the different strains.

Future work would include long-term studies investigating effects of storage conditions to reveal environmental components contributing the loss of SA production in *A. succinogenes*. Additional studies should include transcriptomics to elucidate the underlying shifts in gene expression that cause *A. succinogenes* to change from SA(+) to SA(-). We hypothesize a transcriptomic comparison between SA(+) and SA(-) will reveal differential gene expression related to sigma factors. Previous studies have shown very drastic fermentation profile changes linked to sigma factor switching in bacteria (50, 51). Such patterns revealed in *A. succinogenes* could provide information on how to best maintain SA production in this non-model bacterium long-term. It could also be helpful in pinpointing which regulators are key in how the developed synthetic biology tools are expressed. By identifying potential targets for genetic intervention, *A. succinogenes* could be further developed as a biological platform for bioproduction.

## 3. Conclusion

When working with non-model organisms, the lack of characterization and long-term studies can lead to unexpected challenges. In the example described here, we found that the non-model bacterium *A. succinogenes* demonstrates a shift in fermentation profile and synthetic biology tool expression over time. While this is important information for researchers seeking to develop this particular bacterium, we also present this as a cautionary tale. There is little room for making assumptions when working in non-model organisms. Such organisms with unique traits can be full of surprises and we propose that careful observation over time should be included in studies seeking to add to their characterization. We believe the hurdles to developing non-model bacteria are worth overcoming. As has been seen in other non-model systems, time, effort, and innovative solutions have been able to advance work within organisms of interest (5, 6). By sharing both exciting developments as well as pitfalls, the challenges of non-model systems can be overcome to unlock unique and promising capabilities for use as bio-platforms.

## 4. Materials and Methods

### 4.1 Strain and media recipes

*Actinobacillus succinogenes* 130Z (ATCC 55618) was obtained from the American Type Culture Collection (Manassas, VA) and maintained in tryptic soy broth media (G-Biosciences, St. Louis, MO) with 10 g/L glucose (Fisher Chemical, Hampton, NH) (TSBG hereafter). Seed cultures of *A. succinogenes* were started from 40% glycerol stock in 10 mL medium in 15 mL plastic conical tube and incubated at 37° C, 250 rpm for 14 hours. Fermentation studies were conducted in fermentation medium containing 16.0 g/L yeast extract (Fisher BioReagents, Pittsburgh, PA), 1.0 g/L NaCl (Fisher Chemical), 1.36 g/L NaC_2_H_3_O_2_ (Sigma-Aldric, St. Louis, MO), 0.20 g/L MgCl_2_·6H_2_O (Sigma Life Science, St. Louis, MO), and 0.20 g CaCl_2_·2H_2_O (Fisher Chemical) with D-glucose (Fisher Chemical) and buffered with MgCO_3_ (Acros Organic, Morris Plains, NJ) at 80% the sugar concentration.

### 4.2 Growth study and antibiotic screening procedures

Cultures for growth curve and antibiotic screening procedures were started from seed culture diluted to 0.02 OD_600_ or 2.5% (v/v), respectively, and were grown in 10 g/L TSBG. New tubes were opened at each time point to maintain anaerobic conditions throughout and all measurements were made in biological triplicate. Antibiotics used included kanamycin (Teknova, Hollister, CA), tetracycline (Thermo Scientific, Waltham, MA), ampicillin (Research Products International, Mount Prospect, IL), gentamicin (Fisher Scientific, Waltham, MA), spectinomycin (VWR, Radnor, PA), and chloramphenicol (VWR, Radnor, PA) at concentrations listed in Results and Discussion. Growth repression for each sample was calculated using equation 1. Each sample OD_600_ (*G*_*S*_) was divided by average growth in wild type (*G*_*WTavg*_), converted to a percentage and subtracted from 100 to obtain the percent repression. Growth repression was averaged across three sample replicates for each treatment.

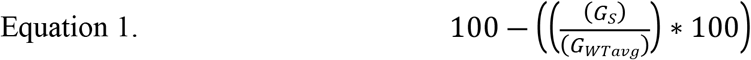

### 4.3 Plasmid construction

For the purpose of plasmid construction, primers were purchased from Eurofins Genomics (Louisville, KY) with no modifications. PCR templates were amplified with Phusion Hot Start II High-Fidelity DNA Polymerase (Thermo Scientific) and amplified products were purified with PureLink Quick PCR Purification Kit (Invitrogen, Carlsbad, CA) or PureLink Gel Extraction Kit (Invitrogen). Sequences of genetic parts, the promoter library, primers used to amplify target sequences, plasmids constructed, and the resulting strains are listed in Supplementary Tables S1-S5 respectively.

Sequences and original sources of key genetic components are listed in supplementary table S1. Hot fusion (52) was used to construct plasmid SSBIO-AS001. Parts included the *E. coli* and *A. succinogenes* origins of replication (originally from the PLGZ920 plasmid (53) generously donated by Dr. Gregg Beckham at the National Renewable Energy Lab in Golden, CO), *gFPuv* (originally from plasmid pBbB7a-GFP (54) purchased from Addgene 35358, Watertown, MA) under the control of P_*lac*_ (originally from *E. coli* strain MG1655 purchased from Yale’s Coli Genetic Stock Center https://cgsc.biology.yale.edu/), and a kanamycin resistance (from the plasmid pBBR1MCS-2 (55) purchased from Addgene 85168). All parts except kanamycin were amplified from a plasmid constructed from a previous project AS-Plac(c)-GFPuv-amp (unpublished, Saha Lab at UNL). The Anderson promoters from the constitutive promoter library (supplementary table S2), were inserted into SSBIO-AS001 by first linearizing the vector, excluding the P_*lac*_ promoter region, using primer pairs that included tails with the promoter library sequences. PCR products were treated with DpnI (Thermo Fisher Scientific, Waltham, MA), gel extracted, phosphorylated with T4 PNK (Thermo Fisher Scientific), and formed into plasmids with T4 DNA ligase (Thermo Fisher Scientific) following the user guide for self-circularization of linear DNA. As the native promoter, P_*pcka*_ (the sequence of which was previously described (21), amplified from A. succiniogenes genomic DNA (extracted with Zymo Research Quick-DNA Fungal/Bacterial Miniprep Kit) was longer, hot fusion was used to combine it with linearized SSBIO-AS001 excluding the P_*lac*_ promoter region. To construct SSBIO-AS002, the promoter variants P_*pcka*_ and BBa-J114 of SSBIO-AS001 were linearized by PCR amplification excluding the *gFPuv* gene region. The flavin binding fluorescent protein gene was synthesized from previously known sequence (56) (Thermo Fisher Scientific) and the plasmid was formed using the hot fusion method. To construct plasmid SSBIO-AS003, the *E. coli* origin of replication, *A. succinogenes* origin of replication, and *gFPuv* under the control of P_*lac*_ as well as the *lacI* gene (originally from *E. coli* strain MG1655) were amplified from the plasmid AS-Plac(i)-GFPuv-amp (unpublished, Saha Lab at UNL) in two pieces. These parts were combined with the previously used kanamycin resistance gene via the hot fusion method. To construct plasmid SSBIO-AS004, SSBIO-AS003, excluding the promoter region, was linearized using primers which contained tails with the sequence for promoter BBa-J23100. Blunt end ligation was used to form the plasmid.

Plasmids expressed in *A. succinogenes* were obtained from cultures of *E. coli* DH10B strain (NEB #C3019 New England Biolabs, Ipswich, MA) using PureLink Quick Plasmid Miniprep Kit (Invitrogen) and were sequence verified. Plasmids were inserted into *A. succinogenes* using a room-temperature electroporation method (Tu et al., 2016) and strains sAS100 – sAS128 (Supplementary Table S5) were frozen at −80° C in 40% glycerol for further analysis.

### 4.4 Expression testing conditions

Expression tests were performed in 10 mL cultures started from 2.5% v/v dilutions of seed cultures and were conducted in biological triplicate. Strains containing the variants of SSBIO-AS001 and SSBIO-AS002 were grown in 10 mL TSBG with 50 µg/mL kanamycin at 37° C, 250 rpm for 8 hours. Strains containing plasmids SSBIO-AS003 and SSBIO-AS004 were grown in 10 ML TSBG with 50 µg/mL kanamycin at 37° C, 250 rpm for 1 hour, IPTG was added in concentrations ranging from 0-5 mM, and cultures were then grown for an additional 8 hours. 500 µL of each culture was pelleted and washed in 500 µL 1x phosphate buffer saline (hereafter PBS) two times. The washed pellets were resuspended in 200 µL 1x PBS and analyzed.

### 4.5 Fluorescence measurements

Measurements were taken using a Molecular Devices SpectraMax i3x Multi-Mode Microplate Detection Platform. Absorbance was measured at 600 nm. GFPuv expression was measured at excitation 395 nm and emission 509 nm after 12 hours of aeration. FbFP expression was measured at excitation 450 nm and emission 495 nm immediately follwing washing. Relative expression for each sample was calculated as shown in equation 2. Fluorescence and absorbance of each sample (*F*_*S*_ and *A*_*S*_ respectively) had average PBS values for fluorescence and absorbance removed (*F*_*Pavg*_ and *A*_*Pavg*_ respectively) and then fluorescence was divided by absorbance. When comparisons were not made to wild type, expression of a wild type control (calculated using fluorescence, *F*_*WT*_, and absorbance, *A*_*WT*_) averaged across 3 replicates was then subtracted to remove background expression. All reported expression values were averaged across 3 replicates.

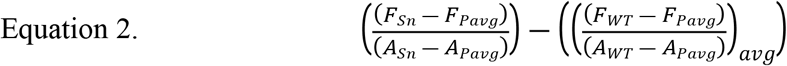

### 4.6 Succinic acid batch fermentation

Seed cultures of *A. succinogenes* were diluted to 2.5% in 100mL of fermentation media in rubber stoppered 125 mL Erlenmeyer flasks in triplicate with between 10 and 50 g/L. The varying sugar concentrations reported are because the data came from comparing a glucose control to another carbon source that varied. Due to making the comparisons across time, we were not able to go back and repeat the studies with equivalent amounts of glucose. The flasks were equipped with check valves to relieve pressure buildup. Cultures were grown at 37 °C and 150 rpm until sampling. Samples were either drawn up through a valve or when cultures were sacrificed. Stoppers were not removed at any point during the fermentation. Growth phase was determined by measuring OD600 of the samples in a spectrophotometer (Thermo Scientific Genesys 10S UV-Vis). Then, cells were pelleted (15 min at 3,000 g) and filtered through 0.45-micron filter into a clean vial.

Concentration of glucose, SA, lactic-, formic- and acetic-acid in the samples were determined by high-performance liquid chromatography (HPLC). A 10 μL cell-free sample was injected into a BioRad chromatographic column (Aminex HPX-87H, 7.8 mm 300 mm, Biorad, USA). The HPLC was equipped with a P680 pump, Dionex, USA and a refractive index detector, RI101 (Shodex, USA). The column temperature was set at 65°C and the eluant used was 0.01 N H2SO4 at a flow rate of 0.6 ml/min.

### 4.7 Genomic sequencing and alignment

Cultures of SA(+) and SA(-) strains were grown to stationary phase and the genomic DNA was extracted (Zymo Research Quick-DNA Fungal/Bacterial Miniprep Kit) for concentrations of 279.1 and 248.9 ng/µl respectively. The samples were sent to the University of Nebraska Medical Center Sequencing Core, Omaha, NE and whole genome sequence data was obtained by Illumina paired end sequencing with read sizes of 150 bases. For each strain, an index was constructed and aligned using Bowtie (57) to the *A. succinogenes* reference genome (GCA_000017245.1). To perform variant calling analysis, Samtools (58) was used to convert from SAM to BAM files and sort them by coordinates. Bcftools was then used to generate the pileups, call SNP and indels, filter high quality loci, and output the variant calling file.

## Supplementary captions

Table S1. Genetics parts and sequences.

Table S2. Promoter library.

Table S3. Primers for plasmid construction.

Table S4. Plasmids used in this study.

Table S5. Strains used in this study.

Figure S1. *A. succinogenes* growth curve with logistic model fit indicating the three growth phases; lag, exponential, and stationary. Error bars represent one standard deviation.

Figure S2. GFPuv expression in *E. coli* strain DH10B with low and high expressing Anderson promoters normalized by absorbance and wild type (Materials and Methods). Error bars represent one standard deviation.

Figure S3. OD_600_ of wild type *A. succinogenes* with varying concentrations of IPTG. Error bars represent one standard deviation.

## Author Information

The authors declare that the research was conducted in the absence of any commercial or financial relationships that could be construed as a potential conflict of interest.

## Author Contributions

DL and CI carried out the experiments. DL wrote the manuscript with support from CI and RS. CI, RS, and MW conceived the original idea. RS supervised the project.

## Acknowledgements

We thank Dr. Gregg Beckham at the National Renewable Energy Lab (Golden, CO) for the generous donation of the *Actinobacillus succinogenes* vector plasmid PLGZ920 used in this study.

